# Comparative analysis of genomic prediction approaches for multiple time-resolved traits in maize

**DOI:** 10.1101/2025.04.22.649925

**Authors:** David Hobby, Robin Lindner, Alain J. Mbebi, Hao Tong, Zoran Nikoloski

**Author notes:** Contributing authors.

## Abstract

Ability to accurately predict multiple growth-related traits over plant developmental trajectories has the potential to revolutionize crop breeding and precision agriculture. Despite increased availability of time-resolved data for multiple traits from high-throughput phenotyping platforms of model plants and crops, genomic prediction is largely applied to a small number of traits, often neglecting their dynamics. Here, we compared and contrasted the performance of MegaLMM and dynamicGP as well as their hybrid variants that can handle high-dimensional temporal data for multi-trait genomic prediction. The comparative analysis made use of time series for 50 geometric, colour, and texture traits in a maize multiparent advanced generation inter-cross (MAGIC) population. The performance of the approaches was assessed using snapshot accuracy and longitudinal accuracy, providing insight into the ability to predict multiple traits at a single time point or the dynamics of individual traits over the considered time domain, respectively.

We found that MegaLMM outperforms dynamicGP in terms of snapshot accuracy, while dynamicGP proved superior in terms of longitudinal accuracy. This study paves the way for careful investigation of factors that affect the capacity to predict dynamics of multiple traits from genetic markers alone.

## 1 Introduction

The plant phenome is a complex outcome shaped by the dynamic interplay of genetic and environmental factors over time. Traditional genomic prediction (GP) models have predominantly focused on predicting traits measured at single time points, providing a snapshot of one or multiple traits in a particular developmental stage for a population of genotypes. However, the advent of high-throughput phenotyping (HTP) technologies has enabled the collection of time-resolved phenotypic data representing hundreds of traits, providing an opportunity to: (*i*) study the trait dynamics over the course of development, (*ii*) better understand the relationships between these traits during the plant life cycle, and (*iii*) use these traits in predicting final agronomically relevant phenotypes, like yield.

Simultaneous prediction of multiple traits, characterizing a part of the entire phenome of a plant, can be achieved by multi-trait GP models [1]. These models make use of genetic and phenotypic correlations among the considered traits, often leading to more accurate predictions compared to traditional single-trait approaches [2–5]. The improved accuracy of multi-trait GP models is largely due to correlations between traits of low heritability to traits of high heritability that can be better predicted from genomic data. However, classical multi-trait genomic prediction (GP) models are computationally demanding, usually addressed by selecting a subset of traits to limit the number of parameters in the model. In addition, with one exception [6], the classical multi-trait GP models have not yet tackled the simultaneous prediction of multiple traits over several time points. Access to and application of such models are of great relevance in practice since they can significantly reduce the efforts for phenotyping of large-scale populations.

Using the premise of the classical GP approach, there are four potential routes to predicting a phenome across time using genomic data: **(1) single-trait models at individual time points**, concerned with predicting each trait at a separate time point. This approach is computationally expensive in terms of the number of models required and fails to exploit information which may be available from other traits or time points. **(2) multi-trait models at a single time point**; here, a model is built for multiple traits jointly at each time point. Although this approach is an improvement over (1) in terms of the number of models trained, it neglects the valuable relationships among traits across different time points. Note that this is distinct from, but methodologically related to, the prediction of the same set of traits and genotypes across multiple years [7]. **(3) single trait across a given range of time points**; this approach predicts one trait over multiple time points within a multi-trait model, but neglects the inter-trait correlations which can contribute considerably to prediction accuracy [6]. Finally, **(4) multi-trait models across a given range of time points**; this is the most comprehensive approach that involves a simultaneous analysis of all traits of interest across all investigated time points. With the ability to capture both inter-trait and inter-time-point correlations, we hypothesize that predictions from approach (4) has the greatest promise to showcase the advantage of genomic prediction to forecast multiple traits at once using genetic markers alone. However, using approach (4) also requires filling a knowledge gap in devising specialized computational and modeling tools that can handle high-dimensional time-resolved data in a genomic prediction setting.

There have been significant advances for single-trait GP models with data from individual time points, and they are widely applied in practice [8]. While several classes of multi-trait GP models have been proposed, facilitating their usage with traits at a single time point, the wider application of these models is precluded largely due to computational demands [1]. The training of single- and multi-trait GP models is performed using different cross-validation (CV) schemes, including CV1 and CV2. Single-trait GP models do not include additional phenotypic data from related traits as “secondary” traits and thus are solely based on genomic data to predict the “focal” trait. On the other hand, multi-trait GP models either can rely solely on genomic data, known as CV1, or can incorporate the same trait, measured in different environments, or different but correlated “secondary” traits, known as CV2, often leading to enhanced prediction accuracy.

A notable exception to the existing multi-trait GP models is the mega-scale linear mixed models (MegaLMM) [9] which can elegantly handle complex, high-dimensional phenotypic data, such as those generated in HTP facilities. MegaLMM is able to support the inclusion of secondary traits to assist in predicting focal traits in a CV2 prediction setup and has proven superior to Genomic Best Linear Unbiased Prediction (GBLUP) in both single- and multi-environment trials for predicting traits like plant height, grain yield, days to silking, and anthesis-silking interval in maize [9]. Further, MegaLMM models applied to data from single field trials outperformed both GBLUP and BayesB in predicting 18 maize stalk quality traits [10]. These models utilized less labor-intensive morphological traits (*i.e*., secondary traits) to predict stalk quality (*i.e*., focal traits), which are more challenging to measure. Although MegaLMM has previously been applied to field data obtained with drone technology for HTP [9], it has not yet been applied to controlled environment HTP, nor to time-series data.

To the best of our knowledge, there is a single approach, termed dynamicGP, that has tackled the problem of simultaneously predicting the dynamics of multiple traits, resulting in multi-trait models across a given range of time points [11]. DynamicGP combines Dynamic Mode Decomposition (DMD) [12] with GP, by applying a DMD algorithm to the trait × time matrix **X** to determine a time-invariant operator (matrix) **A**. This time-invariant operator is subsequently used for the prediction of multiple traits at one time from their value at a preceding time point. This is possible since the operator, **A**, captures the dynamics between traits at consecutive time points. DynamicGP relies on training models based on the widely used single-trait genomic prediction method Ridge-Regression Best Linear Unbiased Prediction (RR-BLUP) [13] to predict the intermediate components **R** and **Φ** of DMD based on Schur decomposition (Schur-DMD) [14] with genomic data [11]. As a result, dynamicGP can predict the dynamics of traits over time, rather than at a single time point. Essentially, this approach predicts the evolution of traits by predicting the matrix **A** for genotypes that were not used in the training set. DynamicGP has two variants: **(1) iterative**, where the true trait values at each time point are used to predict the values at the time point directly following it, and, of more practical value **(2) recursive**, where the trait values at the initial (first) time point are used predict the second, which are then used recursively to predict the traits at all subsequent time points.

One possible drawback of dynamicGP with the RR-BLUP core is that a separate model is required for each element of the predicted matrices. More specifically, predicting the dynamics of *n* traits requires building *r* × *n* + *r* ^2^ RR-BLUP models (see Materials and Methods) of the individual matrix elements, where *r* represents the number of included singular values and vectors. Here, rather than training an RR-BLUP model for each individual element of the Schur-DMD components, we aim to examine the extent to which a MegaLMM model can improve the prediction accuracy of the building blocks of dynamicGP. In addition, we provide a systematic performance comparison of dynamicGP and MegaLMM to predict multiple plant traits measured in a controlled environment HTP facility. To this end, we explore three applications of MegaLMM for time-series analysis: **(1) Replacing the core of dynamicGP**, whereby we substitute the RR-BLUP approach in dynamicGP with MegaLMM to predict the building blocks of dynamicGP, aiming to reduce model complexity and assess the suitability, and any improvement in prediction accuracy. **(2) MegaLMM for full time series prediction**, to which end we apply MegaLMM using the full set of traits at the initial (first) time point as secondary traits and predict the traits over the remaining points in the given time series, representing the same data-usage as recursive dynamicGP with the RR-BLUP core. **(3) MegaLMM in an iterative fashion**, whereby the set of traits at a given time point are predicted using their available measurement at the immediately preceding time point as secondary traits, which serves as a comparison for the iterative version of dynamicGP.

## 2 Materials and Methods

### 2.1 Data generation and pre-processing

A MAGIC maize population of 347 recombinant inbred lines (RILs) and nine founders was phenotyped in three high-throughput experiments at the Leibniz Institute of Plant Genetics and Crop Plant Research in Gatersleben, Germany. Over 347,000 images were captured at multiple time points and processed to derive 498 geometric, colour, and texture traits. A linear mixed model was used to estimate time-point-specific trait heritability and BLUP values as outlined in [11]. Genotyping data for 330 RILs with 70,846 single nucleotide polymorphisms (SNPs) was used for genomic prediction models.

The 498 traits were clustered based on the pairwise Mantel correlations between the genotype by time point matrices for each trait. Each trait was represented by a node in a network, with Mantel correlations above 0.95 represented by an edge between two traits. Clusters of traits were found using modularity clustering as implemented in R package “igraph” [15], from which the trait with the highest mean heritability across all time points was selected as a representative, resulting in 50 representative traits from side, top and combined images. These traits were then normalized and used for subsequent modeling.

### 2.2 DynamicGP implementation

Dynamic Mode Decomposition (DMD) determines a time-invariant best-fit linear operator matrix **A** that transforms a set of measurements, in our case a set of phenotypes, at one time point into the measurements of those same traits at the subsequent time point, *i.e*.

#### 2.2.1 Algorithm 1 (DMD)

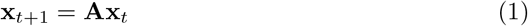

where **x** represents a column vector of *n* traits and **A**, an *n* × *n* matrix. Concatenating the **x** vectors of all time points into matrices, Eq. (1) can be equivalently written as:

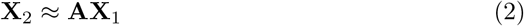

where **X**_1_ and **X**_2_ are *n* × (*T* −1) matrices offset by a single time point (Figure 1a). From **X**_1_ and **X**_2_, the matrix **A** can be directly derived as:

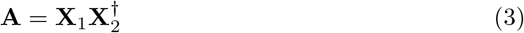

where ^*†*^ indicates the Penrose-Moore pseudoinverse. Algorithm 1, above, allows near-perfect recreation of the training data [11]; however, it is susceptible to noise and requires a large number of models (i.e. *n*^2^ separate models for *n* traits) to be predicted if each matrix element is treated separately. Instead, dynamicGP uses an approximation algorithm termed Schur-DMD [14]:

**Fig. 1.**
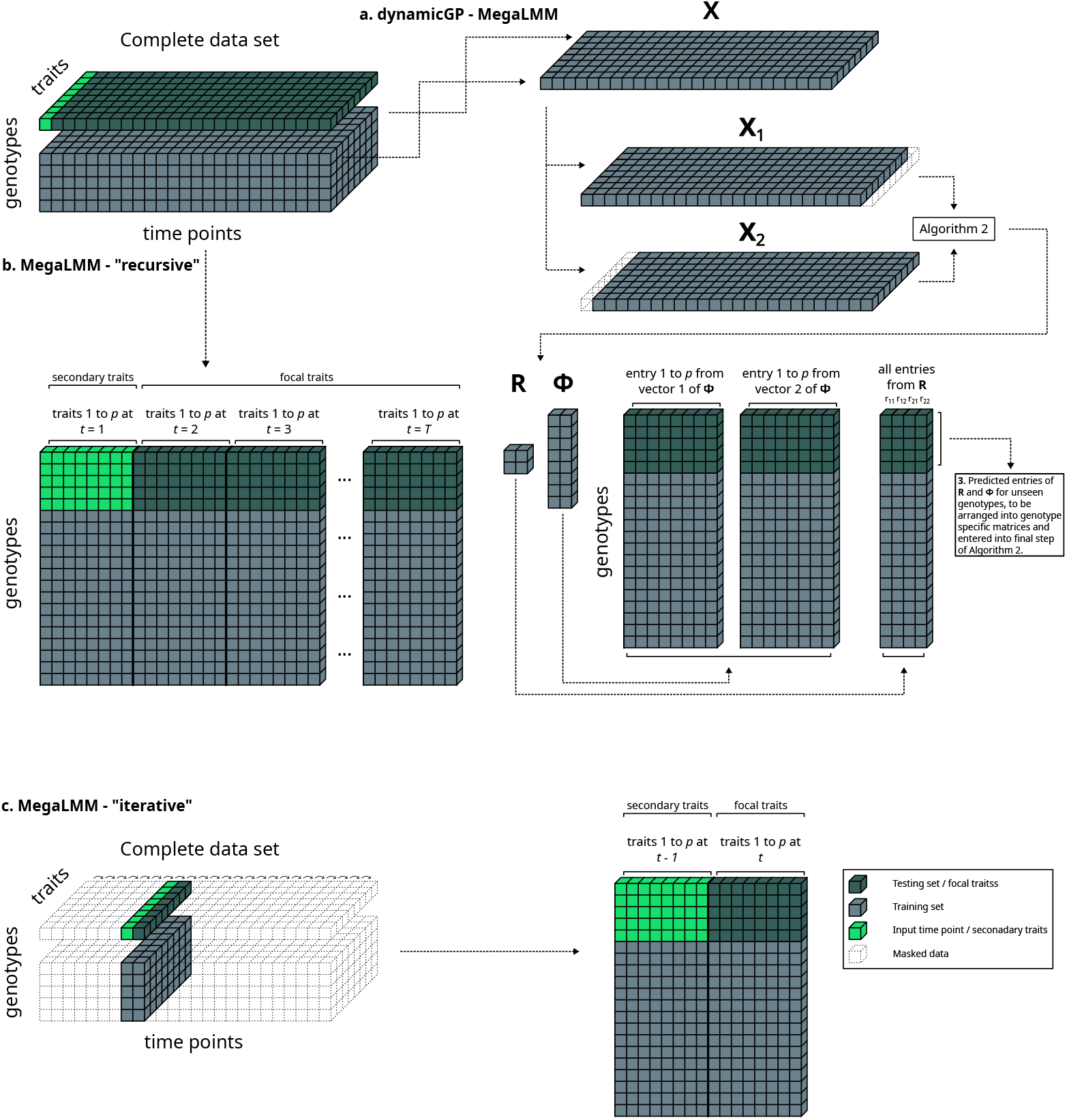
Illustration of the data usage in the compared MegaLMM formulations. **a**. In the first comparison, we used 80% of the data to train dynamicGP, MegaLMM and RR-BLUP models, and employed them to predict the remaining 20% of data. The recursive version of dynamicGP requires the initial (first) time point of the series as input. **b**. MegaLMM models were used to predict all traits and time points simultaneously employing the set of traits at the initial (first) time point as secondary traits. **c**. Iterative MegaLMM models are trained and implemented using the set of traits from the focal time point as training data, along with the set of traits from the immediately preceding time point as secondary traits. This was repeated for each time point, for which a separate model was trained.

#### 2.2.2 Algorithm 2 (Schur-DMD)

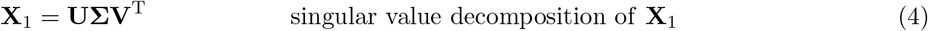

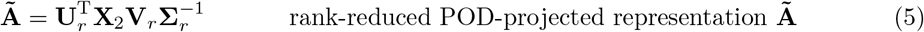

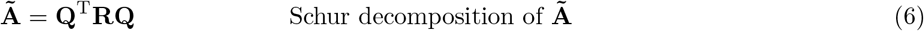

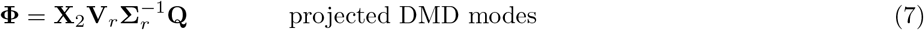

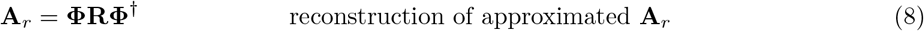

This approximation aims to preserve only the true signals within the data (*i.e*., noise filter). Further, it acts as a dimensionality reduction method, thereby reducing the number of matrix entries required to obtain a predicted **A** matrix from *n*^2^ (*i.e*., Algorithm 1) to *r* × *n* + *r* ^2^ for *n* traits and *r* included singular values from the truncated model in Eq. (5). The parameter *r* was selected by estimating the heritability and prediction accuracy in cross-validation using RR-BLUP models of the elements within the intermediate matrices **U, V**, and **Σ**, corresponding to their singular vector [11]; in our analysis with the data at hand, *r* = 2 was used, as the elements from *r >* 2 had near 0 heritability and prediction accuracy.

The full set of traits at each time point were organized into genotype specific trait × time point **X** matrices, which were then used as inputs to the Schur-DMD algorithm to find the intermediate component matrices **R** (*r* × *r*) and **Φ** (*n* × *r*) (Algorithm 2; Figure 1a). The matrix elements across multiple genotypes from these two matrices are treated as traits in genomic prediction models. DynamicGP initially used separate RR-BLUP models for each trait (denoted here as **dynamicGP-RR-BLUP**); here, we used multi-trait MegaLMM models to predict all the elements of the matrices simultaneously (**dynamicGP-MegaLMM**). Once predictions of the entries of **R** and **Φ** were obtained for testing lines, they were organized into their corresponding matrices, and then used to derive genotype-specific time-invariant operator matrices **A**_*r*_ using Eq. (8), which in turn allow the prediction of traits at subsequent time points from the same traits at a given input time point. We performed 20 iterations of 5-fold cross-validation to predict the components of **R** and **Φ**, which were then employed in Eq. (8) to obtain predictions of **A**_*r*_ for unseen genotypes (Figure 1a).

### 2.3 MegaLMM models

MegaLMM uses a two-level hierarchical model to predict multiple traits simultaneously. First, **Y** ∈ R^(*n* × *t*)^, a matrix of *n* genotypes and *t* traits, is decomposed into lower dimensional *n* × *k* latent trait matrix **F** and *k* × *t* trait loading matrix **Λ**, as well as an *n* × *t* matrix **E** of residuals:

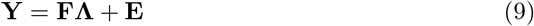

Each of the *k* latent traits in **F** represents the variation of a subset of the original traits and the columns of **F** and **E** are linearly independent and can be modeled as independent linear mixed models, notated as:

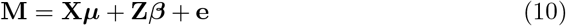

where **M** consists of **F** and **E** concatenated into a single matrix, **X** is an *n* × *p* matrix of fixed effects, ***µ*** is a *p* × 1 vector of fixed effect sizes, **Z** is an *n* × *j* random effects design matrix, ***β*** is a *j* × 1 vector of random effect sizes, and **e** is an *n* × (*k* + *t*) matrix of residuals. By using this two-level approach, MegaLMM avoids performing repeated large matrix inversions typical of multivariate linear mixed models. The number of factors *k* was selected using the recommended *k* = min(*n*/4, *t* /2) [9].

MegaLMM was applied in three ways in the present work: **(1) dynamicGP-MegaLMM**, where we substitute the RR-BLUP core of dynamicGP with MegaLMM to predict the building blocks of dynamicGP (**R** and **Φ**, Figure 1a), **(2) MegaLMM-“recursive”**, where we apply MegaLMM using the full set of traits at the initial (first) time-point as secondary traits and predict the traits over the remaining 24 time points in the given time series - this represents the same data-usage as recursive dynamicGP-RR-BLUP (Figure 1b). **(3) MegaLMM-”iterative”**, where for all time points *t*, the set of traits at time *t* +1 are predicted using their available measurement at time *t* as secondary traits - serving as a comparison for the iterative version of dynamicGP-RR-BLUP (Figure 1c).

### 2.4 Assessment of accuracy

Accuracy of time-series predictions was assessed in two different ways: **(1) snapshot accuracy**, for a trait and time point, Pearson correlation coefficient (PCC) between the true and predicted values of the trait across different genotypes was determined for each time point, resulting in separate prediction accuracies for each trait at each time point, and **(2) longitudinal accuracy**, for a trait and genotype, PCC and Mean squared error (MSE) between the true and predicted values across time of the trait was determined for each genotype. We consider the longitudinal accuracy as the means to assess the accuracy of prediction of a trait’s dynamics. Snapshot accuracies are reported as aggregated accuracies across all traits and CV folds within a time point, while the longitudinal accuracies are aggregated across all traits and lines and CV folds. The accuracy of dynamicGP components was assessed as the mean PCC between the true and predicted values across all CV folds.

## 3 Results

### 3.1 Set-up for comparative analysis of dynamicGP and MegaLMM

The comparative analysis between dynamicGP and MegaLMM utilized 50 geometric, colour, and texture traits measured across 25 time points in 330 maize lines from a MAGIC maize population, resulting in 1250 unique trait-time point pairs (see Materials and Methods). Three distinct MegaLMM approaches were implemented. In the first approach, termed **dynamicGP-MegaLMM**, single-trait RR-BLUP models in dynamicGP were replaced with multi-trait MegaLMM models. Here, the four elements of the **R** matrix (*r* ^2^=4) were predicted simultaneously using a single multi-trait model, while the two vectors of **Φ** (*r* =2), each containing 50 elements corresponding to the 50 traits (Eq. (7), Schur-DMD), were predicted individually using two separate multitrait models. These predictions were then organized into their respective matrices and integrated into the dynamicGP algorithm (Eq. (8), Materials and Methods), requiring three MegaLMM models per cross-validation iteration. Once genotype-specific **A**_*r*_ matrices are calculated they can be employed in iterative or recursive fashion. In the second approach, the **MegaLMM-”recursive”** scenario, a single MegaLMM model was trained using all 50 traits at *t* =1 as secondary traits to predict these traits across time points 2 to 25, yielding predictions for 1200 trait-time-point pairs (Figure 1b). In the third approach, the **MegaLMM-”iterative”** scenario, multi-trait MegaLMM models were trained iteratively, using all traits at one time point (*e.g*., *t*) as secondary traits to predict traits at the next time point (*t* +1), repeating this process across the time series from *t* =1 to *T* −1 (Figure 1c), requiring a separate model for each time point.

### 3.2 MegaLMM fails to accurately predict dynamicGP building blocks

In this study, dynamicGP was applied to multi-trait time-series data comprising 50 traits measured across 25 time points. Individual models were constructed to predict the elements of the **R** and **Φ** matrices, which were then arranged into their corresponding matrices to calculate genotype-specific **A**_*r*_ matrices for longitudinal predictions (see Materials and Methods). While the original dynamicGP implementation utilized *r* × *n* + *r* ^2^ RR-BLUP models, these were replaced with three MegaLMM models in the present study (see Materials and Methods). Using MegaLMM to predict the **R** and **Φ** components, which were subsequently integrated into dynamicGP, resulted in a prediction accuracy of 0 across all time points for both recursive and iterative configurations (Figure 2). Further analysis of the prediction accuracy for the building blocks of dynamicGP revealed that MegaLMM models yielded near-zero mean accuracies across 20 iterations of 5-fold cross-validation for most elements of the matrix components (Figure 3a). Notably, the **Q** and **Φ** components exhibited mean prediction accuracies of 0.05 and 0.07, respectively, although these values remained lower than those achieved with RR-BLUP models for all components (Figure 3a). In addition, the inter-element correlations in dynamicGP components were assessed, showing a mean correlation near 0 for most components, with non-zero mean correlations observed for **Σ, R**, and **Φ** (Figure 3b). This finding can explain the poor performance of the hybrid variant of dynamicGP with MegaLMM models for its building components.

**Fig. 2.**
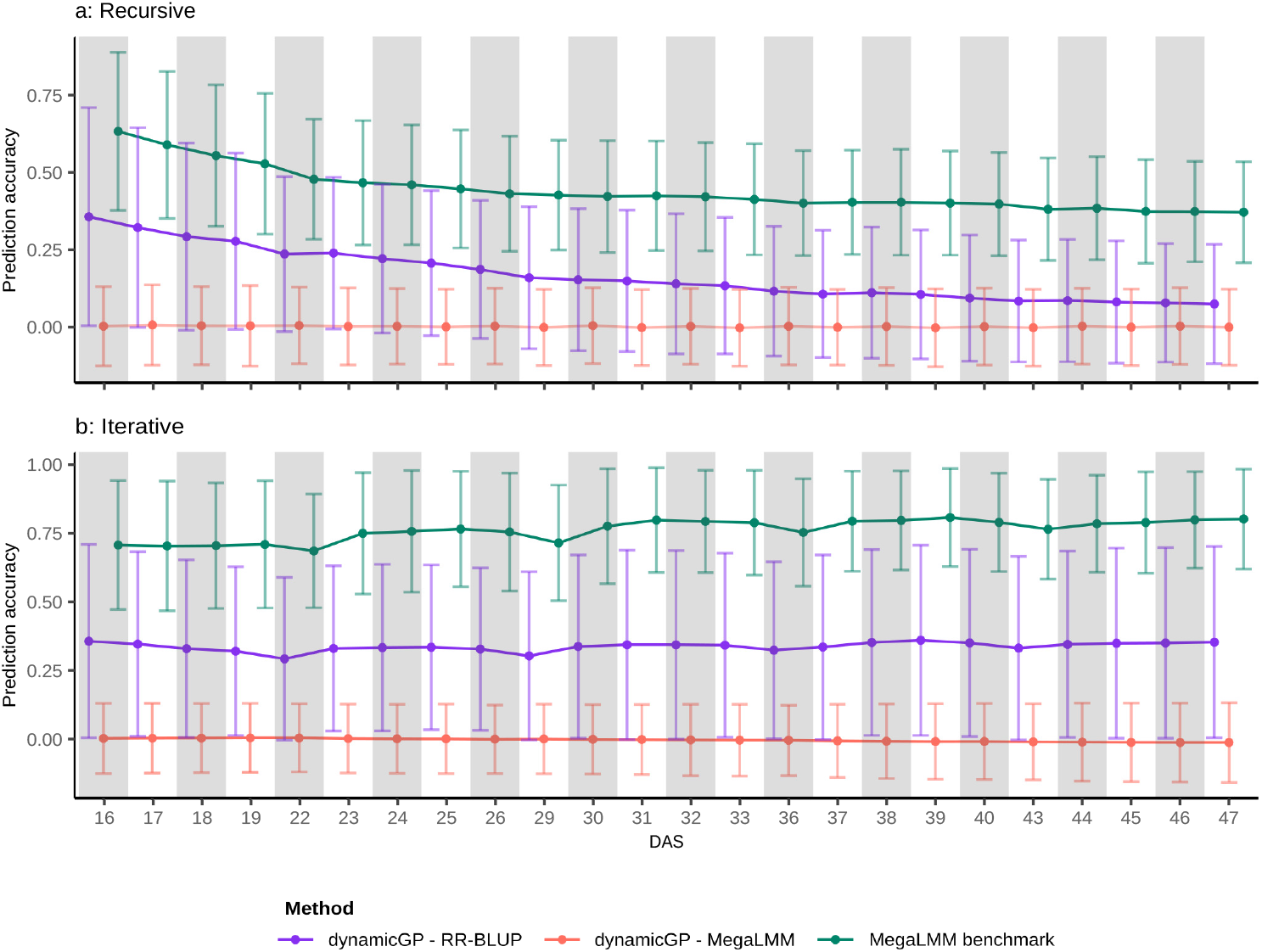
Comparative analysis of MegaLMM and dynamicGP variants based on snapshot accuracy. **a**. Given the measurements of traits at the initial time point to predict the full multi-trait time-series, MegaLMM models outperformed the recursive dynamicGP with RR-BLUP across the full time-series. **b**. Given the measurements of traits at a time point as secondary traits to assist the prediction of these traits in the subsequent time point, MegaLMM models outperformed the iterative dynamicGP-RR-BLUP across the full time-series. In both scenarios dynamicGP-MegaLMM failed to yield any non-zero prediction accuracies.

**Fig. 3.**
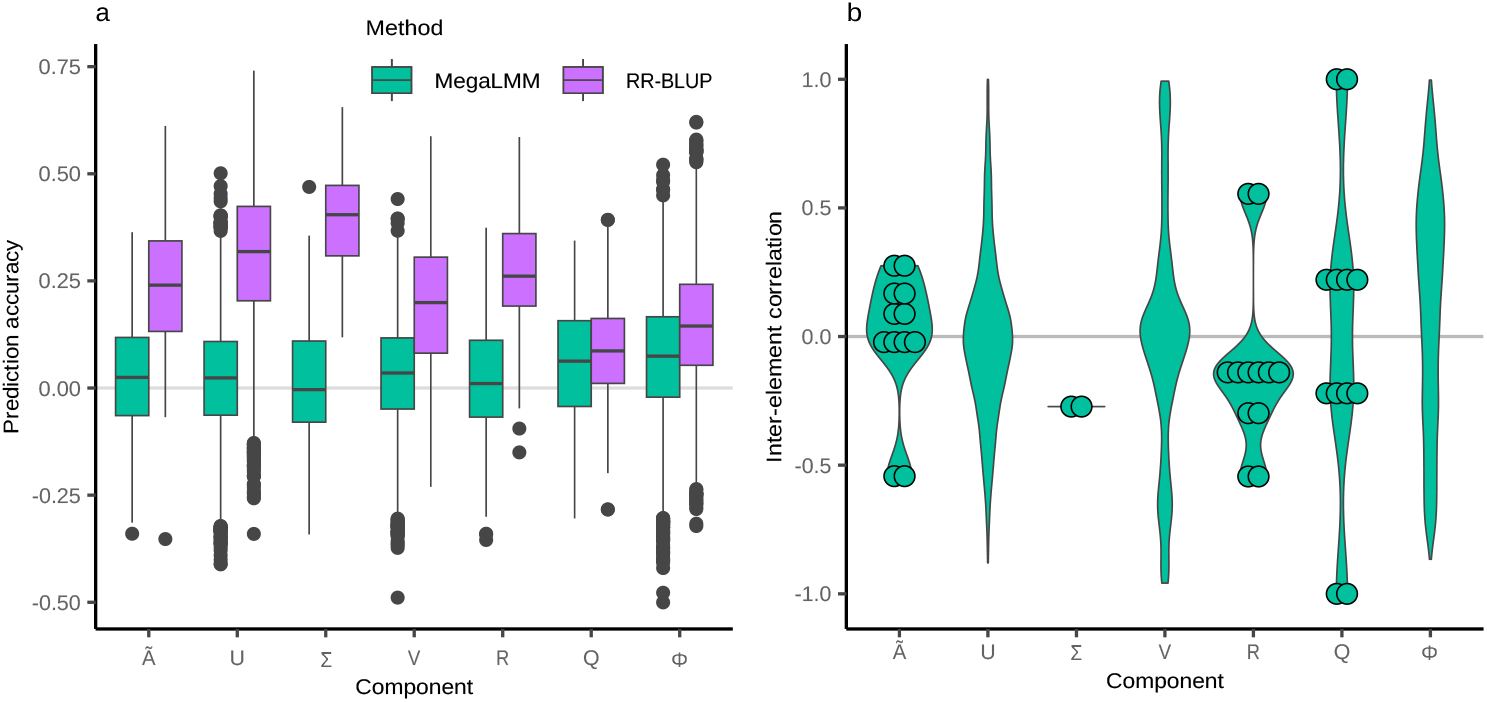
Inter-element correlations in the building blocks of dynamicGP affects the performance of its MegaLMM hybrid variant. **a**. The mean prediction accuracies of the majority of intermediate components of dynamicGP with MegaLMM were near zero; we note that only the components of **Q** and **Φ** had non-zero prediction accuracies. In contrast, RR-BLUP yielded non-zero prediction accuracies for all components. **b**. The mean inter-element correlation within all the intermediate components except for **Σ, R** and **Φ** was zero. Points represent inter-element correlations for matrices with four or fewer elements.

### 3.3 MegaLMM applied directly to time-series data outperforms dynamicGP-RR-BLUP in snapshot accuracy

MegaLMM was applied directly to the time-series data in two configurations in addition to its use within dynamicGP. In the first configuration (complete time-series prediction), 50 traits at *t* =1 were used as secondary traits to predict the remaining 1200 unique trait-time point pairs (50 traits across 24 time points) from genomic data, aligning with the data usage of recursive dynamicGP. In the second configuration (iterative time-series prediction), 50 traits at a given time point (*e.g*., *t* =1) were used as secondary traits to predict the same traits at the next time point (*e.g*., *t* =2), with this process repeated sequentially using separate models for each transition (*e.g*., *t* =2 to *t* =3), utilizing less data than iterative dynamicGP by including only the focal and immediately preceding time points.

For the complete time-series configuration, MegaLMM-”recursive” - MegaLMM outperformed recursive dynamicGP-RR-BLUP by approximately 265% in snapshot accuracy across the full time series (Figure 2a). Prediction accuracies decayed over time, with recursive dynamicGP-RR-BLUP starting at 0.35 ± 0.35 at *t* =16 and dropping to 0.07 ± 0.19 at *t* =47, while MegaLMM began at 0.63 ± 0.26 at *t* =16 and ended at 0.37 ± 0.16 at *t* =47 (Figure 2a). MegaLMM-”iterative” outperformed recursive dynamicGP-RR-BLUP by approximately 226% across the time series (Figure 2b), achieving a mean prediction accuracy of 0.76 ± 0.20 compared to 0.34 ± 0.33 for dynamicGP-RR-BLUP. In this configuration, prediction accuracy dipped noticeably at every fifth time point (Figure 2b), and a slight increase in accuracy was observed toward the end of the time series.

### 3.4 DynamicGP more accurately reproduces trait dynamics

To assess the ability of dynamicGP and MegaLMM to predict trait dynamics beyond snapshot accuracies, longitudinal prediction accuracy was evaluated using Pearson correlations between predicted and true trait values across time points for each trait and genotype, across 20 iterations of 5-fold cross-validation. Iterative dynamicGP-RR-BLUP achieved the highest mean longitudinal correlation of 0.32 ± 0.35 across all genotype-trait combinations, followed by recursive MegaLMM with 0.17 ± 0.50 (Figure 4a). Recursive dynamicGP-RR-BLUP yielded a mean correlation of 0.12 ± 0.46, while recursive MegaLMM had a mean of 0.00 ± 0.53 (Figure 4a). Despite higher mean accuracies, iterative and recursive dynamicGP-RR-BLUP produced fewer significant correlations after Bonferroni correction (13,096 and 10,144, respectively) compared to iterative and recursive MegaLMM (20,767 and 20,939, respectively). Log-adjusted mean squared errors (MSE) were lower for dynamicGP-RR-BLUP, with means of −3.41 ± 1.01 (iterative) and −3.13 ± 0.97 (recursive), compared to −0.29 ± 3.51 (iterative MegaLMM) and −0.61 ± 3.34 (recursive MegaLMM), with MegaLMM MSEs showing high positive skew (Figure 4b).

**Fig. 4.**
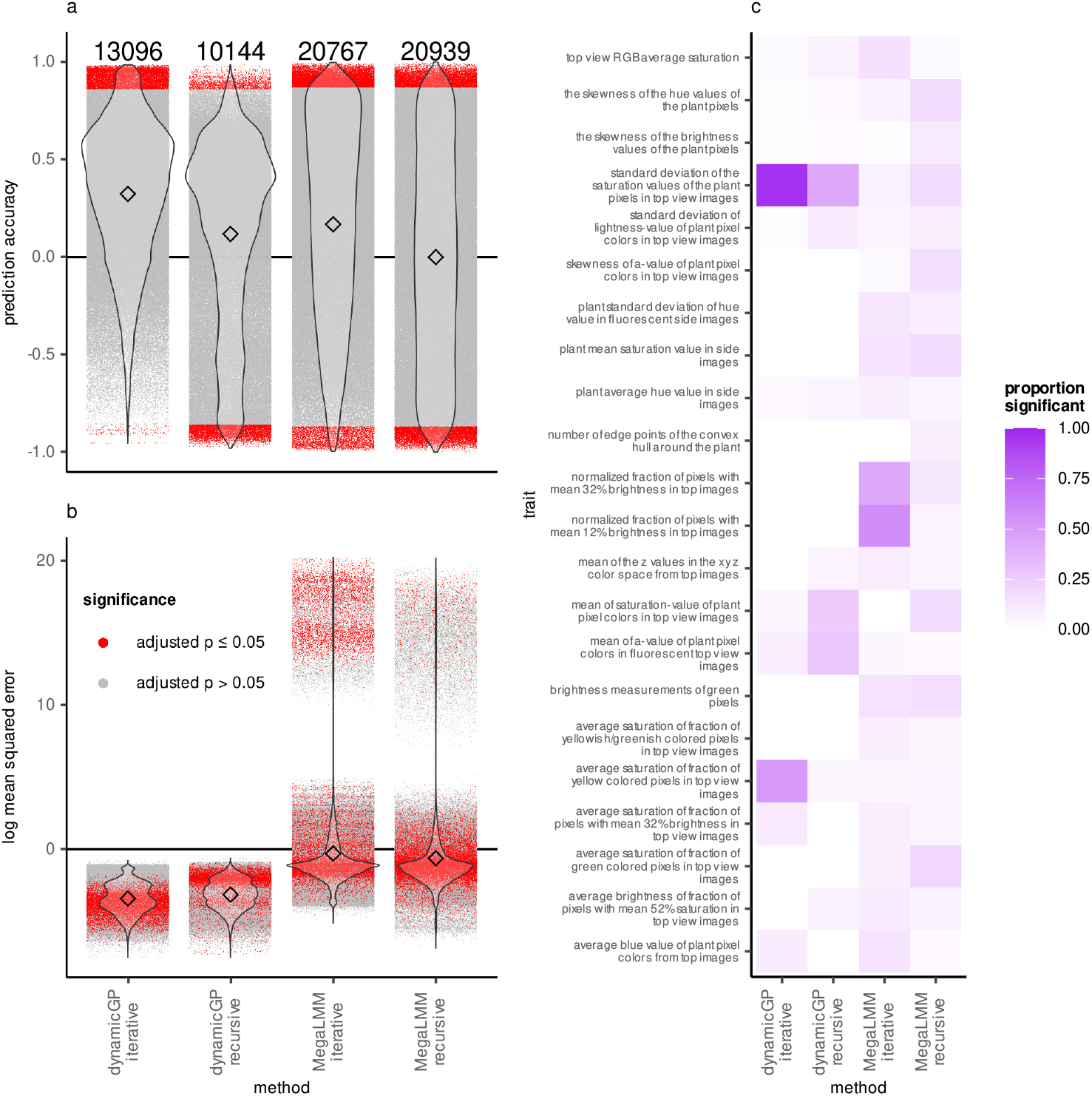
DynamicGP outperforms MegaLMM in terms of longitudinal accuracy. **a**. Accuracy of predicted trait dynamics along time series aggregated across all iterations, traits, and genotypes for MegaLMM as well as both iterative and recursive variants of dynamicGP. Accuracy was assessed as the Pearson correlation between true and predicted values. The numbers along the top indicate the number of significant correlations after Bonferroni correction of *p*-values. **b**. Mean squared error of predicted trait dynamics along time series aggregated across all iterations, traits, and genotypes for dynamicGP in recursive and iterative variants as well as the MegaLMM comparisons. Within each method there are a total of 330,000 tests performed, corresponding to the total number of combinations of 50 traits over 330 genotypes and 20 cross-validation, with each indicated by a point. Red points indicate significant correlations across both panels after Bonferroni correction of *p*-values. **c**. Proportion of significant predictions for traits using each of the four methods. For an extended heatmap containing all traits see Supplementary Figure 1.

The proportion of significant correlations was calculated for each trait across 6,600 predictions (330 genotypes × 20 iterations). For iterative MegaLMM, two traits — *normalized fraction of pixels with mean 12% brightness in top images* and *normalized fraction of pixels with mean 32% brightness in top images* — showed significant correlations in 58% and 45% of tests, respectively, while other traits averaged 7% (Figure 4c). For iterative dynamicGP-RR-BLUP, two traits - *standard deviation of saturation values of plant pixels in top view images* and average saturation of the yellow-colored pixel fraction in top view images - exhibited significant correlations in 95% and 50% of tests, respectively, with the remaining traits averaging 4% (Figure 4c). In contrast, recursive dynamicGP-RR-BLUP showed a mean of 3% significant tests, with the standard deviation of saturation values also having the highest proportion. MegaLMM-”recursive” averaged 7% significant tests across traits (Figure 4c).

## 4 Discussion

The three MegaLMM implementations provided distinct frameworks for comparison with dynamicGP-RR-BLUP, each providing different means for using genomic data. The dynamicGP-MegaLMM approach extended the original dynamicGP by incorporating multi-trait genomic predictions for **R** and **Φ** components, enabling simultaneous modeling of multiple elements. This hybrid method contrasts with the MegaLMM-“recursive” scenario, which utilized a single model to predict 1200 trait-time-point pairs based solely on initial time point data (*t* =1), offering a comparison to the recursive dynamicGP-RR-BLUP setup. In comparison, the MegaLMM-”iterative” scenario used step-wise prediction, training separate models for each time point, which mirrors the iterative dynamicGP-RR-BLUP configuration, but takes advantage of inter-trait correlations within and between individual time points. The MegaLMM-”recursive” approach stands out for its ability to handle the full time series in a single model, potentially reducing computational overhead compared to the MegaLMM-”iterative” approach, which requires 24 distinct models for the 25 time points. These configurations highlighted trade-offs between model complexity, predictive scope, and alignment with dynamicGP-RR-BLUP’s recursive and iterative frameworks, providing a basis for evaluating their performance against dynamicGP-RR-BLUP (see Materials and Methods).

The replacement of RR-BLUP models with three MegaLMM models in dynamicGP-MegaLMM aimed to use the increase in accuracy of multi-trait prediction, but yielded a mean prediction accuracy of zero across all time points in both recursive and iterative configurations (Figure 2). This indicating that MegaLMM as employed here was not a suitable replacement for RR-BLUP (Figure 2). This poor performance prompted an investigation into MegaLMM’s prediction accuracy of the elements of the intermediate matrices **R** and **Φ** as well as the other matrices from the Schur-DMD algorithm, which revealed generally low accuracies, with only **Q** and **Φ** reaching modest values of 0.05 and 0.07, respectively, which are lower than the RR-BLUP benchmarks (Figure 3a). Given that MegaLMM exploits inter-trait covariance to enhance prediction accuracy over single-trait methods, the near-zero mean correlations among most dynamicGP component elements (Figure 3b) likely undermined its effectiveness. Although non-zero correlations were observed for **Σ, R**, and **Φ** (Figure 3b), these were insufficient to provide a predictive advantage. Consequently, the lack of strong inter-element correlations may not only have negated the benefits of a multi-trait approach but also degraded MegaLMM’s performance, driving prediction accuracies toward 0. These findings indicated that the structures of the analyzed matrices do not align with MegaLMM’s strengths, reinforcing the suitability of RR-BLUP for predicting dynamicGP components.

The direct application of MegaLMM to time-series data revealed distinct performance advantages over recursive dynamicGP-RR-BLUP in both configurations. Using MegaLMM-”recursive” model, we found a 265% improvement in accuracy across all traits and time points (Figure 2a). This demonstrates its ability to extract meaningful information from the initial trait data (*t* =1), used as secondary traits, for robust long-term predictions, unlike dynamicGP-RR-BLUP, where error propagation, accumulating from each prediction step, likely drove the accuracy decline (from 0.35 to 0.07). The decay in accuracy for MegaLMM (0.63 to 0.37) may stem from decreasing trait autocorrelation as temporal distance increases, reducing the predictive power of early time points compared to later ones. The gain in accuracy of the MegaLMM-“iterative” model over dynamicGP-RR-BLUP (mean 0.76 vs. 0.34) reflects its effective use of pairwise time-point data, although it uses less data than iterative dynamicGP-RR-BLUP, suggesting that immediate trait relationships are sufficient for accurate predictions (Figure 2b). The observed accuracy dips at every fifth time point, aligning well with a two-day measurement gap after each five-day interval, causing weakened trait correlations compared to the 24-hour gaps between all other sequential time points; this observation in part explains the periodic reductions in accuracy. The slight accuracy increase late in the time series could indicate that genetic differences become more pronounced in mature plants, enhancing MegaLMM’s ability to distinguish trait variations compared to earlier stages. These results suggested the superiority of MegaLMM in direct time-series applications, particularly when error propagation or data gaps impact the performance of dynamicGP.

While snapshot accuracies provide a basis for ranking and selecting genotypes, they do not capture the ability to predict temporal trait trajectories, as high Pearson correlations at individual time points may not accurately reflect the shape of the trait trajectories. Longitudinal analysis revealed that iterative dynamicGP-RR-BLUP outperformed other methods with a mean correlation of 0.32, suggesting better accuracy in tracking trait dynamics over time, despite fewer significant correlations (13,096) compared to MegaLMM (up to 20,939). The near-zero accuracy of MegaLMM-”recursive” indicates a failure to capture longitudinal patterns, possibly due to its reliance on broad trait relationships that weaken over time, whereas dynamicGP-RR-BLUP’s lower but positive accuracies (0.12–0.32) and reduced MSEs (−3.41 and −3.13 vs. −0.29 and −0.61) suggest greater ability to predict trajectories of multiple traits (Figure 4a & b). The skewed MSEs of MegaLMM further highlight its inconsistency in this context.

Trait-specific analysis revealed that the superior performance of dynamicGP-RR-BLUP may stem from exceptional accuracy in a few traits (*e.g*., 95% and 50% significant tests for *standard deviation of the saturation values of the plant pixels in top view images* and *average saturation of fraction of yellow colored pixels in top view images*, respectively), while most traits were poorly predicted (mean 3% and 4% for iterative and recursive, respectively), inflating the overall average (Figure 4c). In contrast, MegaLMM’s higher number of significant correlations (*e.g*., 58% and 45% for *normalized fraction of pixels with mean 12% brightness in top images* and *normalized fraction of pixels with mean 32% brightness in top images*, respectively) indicates broader but less precise predictive success across traits, averaging 7%. The inverse relationship between iterative and recursive dynamicGP-RR-BLUP accuracies for certain traits (Supplementary Figure 2) suggests configuration-specific strengths, potentially tied to trait dynamics or data structure. For instance, *normalized fraction of pixels with mean 12% brightness in top images*, poorly predicted by recursive dynamicGP, was well predicted by MegaLMM-iterative, suggesting dependence of predictability on a particular model-trait combination. It is worth noting that *normalized fraction of pixels with mean 12% brightness in top images* was also found to be the trait with the lowest mean prediction accuracy across all time points with recursive dynamicGP-RR-BLUP [11]. These findings underscore the ability of dynamicGP-RR-BLUP to predict trajectories for selected traits, while MegaLMM offers wider but less accurate longitudinal prediction performance. Our findings reflect the methodological differences underpinning the models and their ability to capture temporal and trait-specific variation across genotypes.

## 5 Conclusion

Longitudinal predictions from genomic data are agronomically important as they allow for a shortening of the breeding cycle and resources for phenotyping, which represent significant bottlenecks in modern breeding. Typically, in time-dependent genomic prediction studies, accuracy is assessed using the snapshot method quantified by the Pearson correlation between measured and predicted values separately at each time point. In the present study we additionally examined longitudinal accuracy - the Pearson correlation between the true and predicted trait development trajectories over multiple time points. We compared the performance of a powerful multi-trait genomic prediction method, MegaLMM, with that of a recently introduced method, dynamicGP, in predicting a multi-trait time-series of 50 traits. The performance of the two approaches was assessed using both snapshot and longitudinal accuracies. We found that while MegaLMM outperformed dynamicGP in terms of snapshot accuracy across all time points, it produced predictions which were not representative of the true dynamics of the traits across time. Our findings were observed in a single data set from a single species, and further research is required the generality of these observations on other data sets. Furthermore, our study points at the need for additional methodological developments to tackle the challenging problem of multi-trait temporal prediction from genetic markers.

## Supplementary information

Supplementary Figures 1 & 2 are found in file “Supplementary Figures.pdf”.

## Declarations

### Funding

Part of this research was funded by the Horizon Europe research and innovation program, project BOLERO (Breeding for coffee and cocoa root resilience in low-input farming systems based on improved rootstock, HORIZON-CL6-2021-BIODIV-01-13), under grant agreement ID: 101060393 (to Z.N.). The project was also supported by the European Union’s Horizon 2020 research and innovation program grant 862201 (to Z.N.).

### Conflict of interest

The authors declare no competing interests.

### Data availability

Genotypic and phenotypic data for maize are available at 10.5281/zenodo.14959484.

### Code availability

All code that was used to generate the results of this study is available at GitHub https://github.com/dobby978/dynamicGP.

### Author contribution

D.H. analyzed data, implemented and tested the approach, performed research. A.M., H.T., and R. L. performed research and contributed to the development and refinement of the approach. Z.N. designed research, supervised research, acquired funding, and contributed to the development of the approach. D.H and Z.N. wrote the first version of the manuscript. All authors contributed to revising and finalizing the paper.

## Notes

https://doi.org/10.5281/zenodo.14959483

https://github.com/dobby978/dynamicGP

### Competing Interest Statement

The authors have declared no competing interest.

