## Supplementary Information for "Comparative analysis of genomic prediction approaches for multiple time-resolved traits in maize"

### **Supplementary Figures**

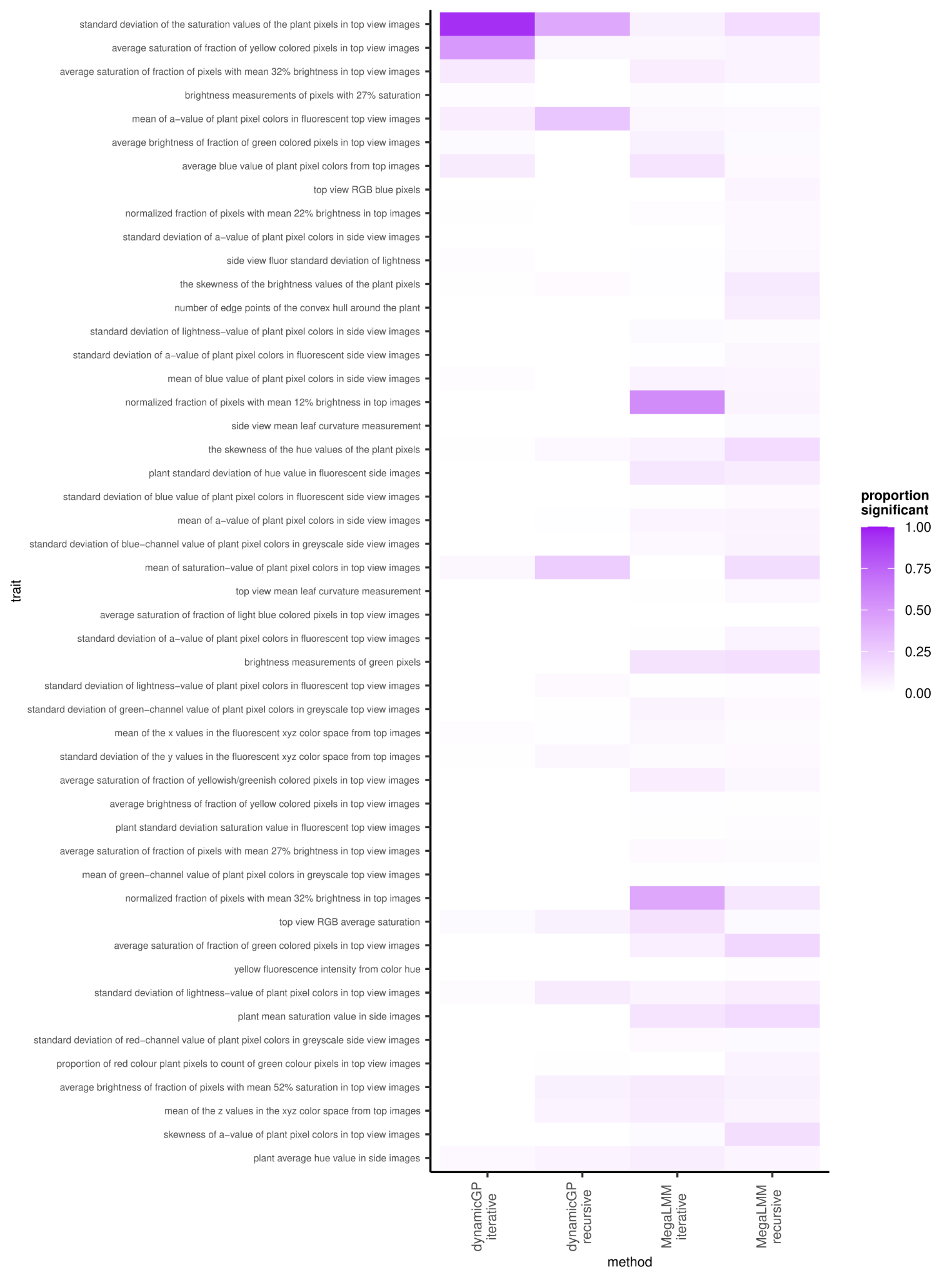

**Supplementary Fig. 1 Proportion of significant predictions for all traits. The** longitudinal accuracy of each trait was assessed 6600 times (330 genotypes × 20 cross-validations). This heatmap shows the proportion of those tests which were significant with *p <* 0.05 after Bonferroni correction.

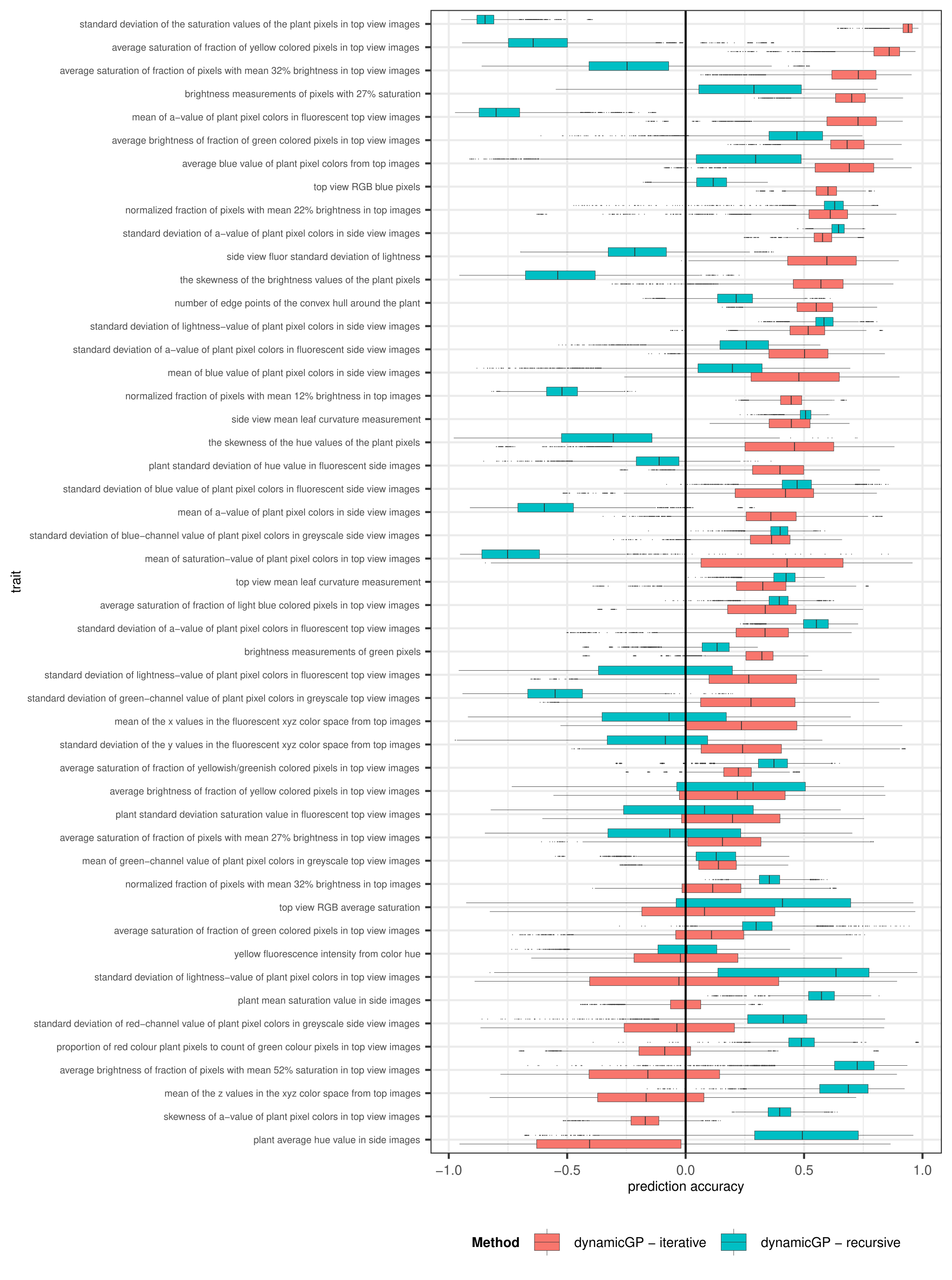

**Supplementary Fig. 2 High accuracy in iterative dynamicGP does not imply high accuracy in recursive.** Per trait accuracy of predicted trait dynamics along time series aggregated across all iterations and lines for dynamicGP in recursive and iterative formats. Accuracy assessed as the Pearson correlation between true and predicted values.

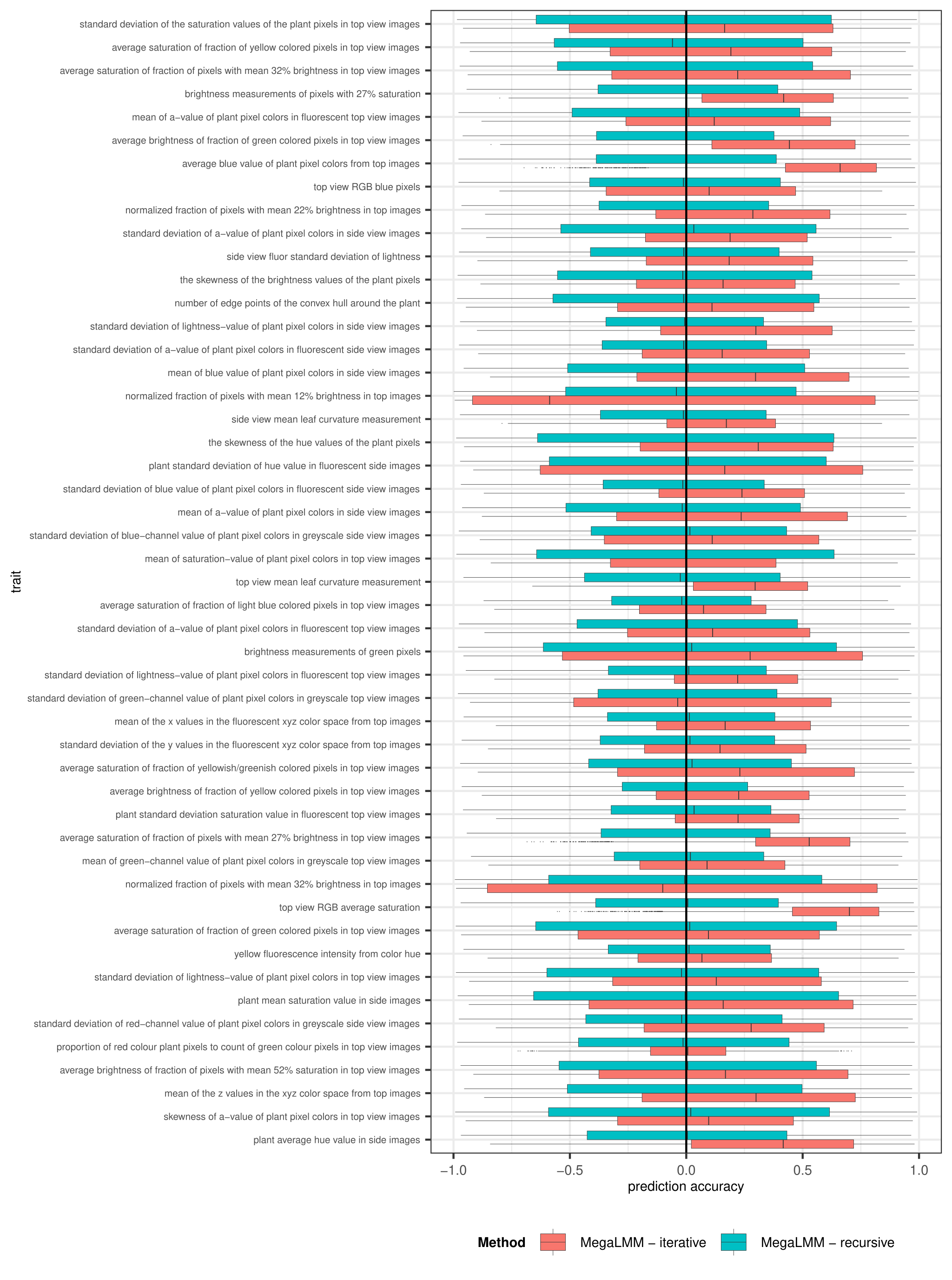

**Supplementary Fig. 3 MegaLMM-iterative outperforms MegaLMM-recursive in terms of longitudinal accuracy.** Per trait accuracy of predicted trait dynamics along time series aggregated across all iterations and lines for MegaLMM in recursive and iterative formats. Accuracy assessed as the Pearson correlation between true and predicted values.
